# Lipidation of a bacterial effector is critical for bacterial evasion of host-defense

**DOI:** 10.1101/2020.11.05.369652

**Authors:** Natalia Cattelan, Hongjiao Yu, Kornelia Przybyszewska, Rosa Angela Colamarino, Massimiliano Baldassarre, Stefania Spanò

## Abstract

The Rab32 antimicrobial pathway has been shown to restrict *Salmonella* Typhi, in mouse macrophages. The broad-host pathogen *Salmonella* Typhimurium however has evolved a strategy to evade the Rab32 antimicrobial pathway, via its effector protein GtgE. GtgE is a cysteine protease that specifically mediates the cleavage and inactivation of Rab32. Here we show that GtgE association and targeting to membranes is critical for its efficient proteolytic activity. The C-terminus of GtgE contains a CaaX motif, which can be post-translationally modified by the host’s prenylation machinery. Using a combination of confocal microscopy and subcellular fractionation we show that a cysteine in the CaaX motif is crucial for GtgE membrane targeting and, more importantly, GtgE localization to the *Salmonella*-containing vacuole. We also demonstrated that prenylation of CaaX is important for an effective and fast Rab32 cleavage, which in turn helps *Salmonella* to successfully survive in macrophages and establish an *in vivo* infection in mice. Our findings shed light on the importance of a host mediated post-translational modification that targets GtgE to the membranes where it can efficiently cleave and inactivate Rab32, leading to a better *Salmonella* survival in macrophages.

**Author summary:** *Salmonella* species includes a large group of bacteria that cause disease in different hosts. While some serovars are host generalists, others are restricted to humans. This is the case of *Salmonella* Typhi, responsible for Typhoid fever, a disease that affects millions globally. We have previously discovered an antimicrobial activity in macrophages that is controlled by Rab32. While the broad-host bacterium *Salmonella* Typhimurium effectively counteracts this mechanism through the delivery of two effectors, GtgE and SopD2, *Salmonella* Typhi does not express those effectors and cannot survive in mouse macrophages. In this article, we demonstrate how *Salmonella* Typhimurium exploits a host machinery to modify GtgE. We show that this host mediated modification is important for GtgE intracellular localization and effective Rab32 targeting, resulting in both a better intracellular survival and infection in vivo.

## Introduction

*Salmonella enterica* includes numerous serovars that cause a broad range of diseases. Common gastrointestinal diseases are caused by broad-host serovars such as *Salmonella* Typhimurium (1) whereas an invasive systemic life-threatening disease, known as Typhoid fever, is associated with human-restricted serovars *Salmonella* Typhi and *Salmonella* Paratyphi (2). An important trait of the *Salmonella* infectious process is the intracellular survival of the bacterium in macrophages, dendritic cells and epithelial cells by establishing a replicative niche either inside of a vacuole known as a *Salmonella*-containing vacuole (SCV) or in the cytosol of the cells (3). In order to successfully do this, the bacterium is equipped with multiple Type III Secretion Systems (TTSS) and a battery of effectors (4).

In some cases, bacterial effectors can be targeted to subcellular compartments by host-mediated post-translational modifications, such as ubiquitination, lipidation or phospholipid binding (5). Consequently, an effector that is delivered to the cytoplasm in low concentration can be accurately targeted to its site of action, increase its effective concentration and ensure engagement with its interactor. One such mechanism is prenylation; a post-translational modification in which an isoprenoid (farnesyl or geranylgeranyl) is irreversibly added to a Cys residue in a CaaX conserved motif at the C-terminus of a protein (6). This motif is recognized and targeted by protein geranylgeranyl transferase I (PGGTI), protein farnesyl transferase (PFT) or Rab geranylgeranyl transferase (RGGT) enzymes to introduce an isoprenoid to the cysteine (7). Once prenylated, the protein is transported to the cytosolic side of the Endoplasmic Reticulum membrane where the aaX C-terminal peptide will be cleaved, and the prenylated Cys will be methylated. Consequently, a hydrophobic domain is added to the protein, which results in its localization to an intracellular membrane compartment (reviewed in (8)).

In this work we studied the prenylation of GtgE, a critical SPI-1/2 TTSS *Salmonella* effector, and its importance for the evasion of the Rab32 antimicrobial pathway in mice. Rab32 is a Rab-GTPase involved in lysosome-related organelles trafficking (9). Like any other Rab-GTPase, Rab32 functions as binary molecular switch, with two states: an inactive state, when it is bound to GDP, and an active state complexed to GTP (10). Cycling between these two states is regulated by the action of guanidine nucleotide exchange factors (GEF), which activate GTPases exchanging GDP by GTP, and GTPase activating proteins (GAP), which stimulate GTP hydrolysis (10). Rab-proteins attach to membranes after addition of one or two geranylgeranyl lipids at their C-terminus. While active Rab-proteins will remain associated with membranes, inactive Rabs can be targeted by GDP-dissociation factor (GDI), solubilizing them into the cytosol (10).

Recruitment of Rab32 to the SCV in infected murine macrophages was observed to be associated with bacterial killing (11-13). It is hypothesized that Rab32 is involved in delivering an antimicrobial cargo to the SCV, which Chen et al. recently showed could be itaconic acid (14). GtgE is a cysteine protease that plays a role in the inactivation of the Rab32 antimicrobial pathway by cleaving Rab32 (11). GtgE acts cooperatively with SopD2, a SPI-2 TTSS effector that generates an inactive Rab32-GDP bound form by its GAP function (12). When *S*. Typhimurium infects a mouse macrophage, Rab32 is not recruited to the SCV due to the activity of both GtgE and SopD2 functions, and the bacterium is able to survive intracellularly (12). Although the roles of GtgE and SopD2 roles are redundant, both are needed for intracellular survival and full virulence of *Salmonella* (11, 12). *S*. Typhi however, does not express either GtgE or SopD2; as result, Rab32 is recruited to the *S*. Typhi-containing vacuole and the bacterium is not able to survive in murine macrophages (11).

In this paper, we show that host-mediated prenylation of GtgE after infection is crucial for its activity towards Rab32-dependent antimicrobial activity. Moreover, a replacement of the Cysteine residue in the CaaX motif of GtgE with a Serine, prevents prenylation and affects its intracellular localization. This indicates that GtgE may need to be prenylated in order to be directed to membranous compartments. We also found that post-translational lipidation affects the ability of GtgE to cleave Rab32 in infected bone-marrow derived macrophages (BMDM). Finally, we observed that mutation of the CaaX sequence impairs the ability of *Salmonella* to survive in murine macrophages and in mice, suggesting that targeting of GtgE to the membranes is important for *in vivo* pathogenicity.

## Results

### GtgE localization is dependent on prenylation

We have previously shown that the ability of *Salmonella enterica* Typhimurium (*S*. Typhimurium) to infect mice depends, at least in part, on targeting the Rab32 GTPase in macrophages through the delivery of GtgE and SopD2 effectors (11, 12). Moreover, a *S*. Typhimurium mutant defective for both these effectors is virtually avirulent in wild type mice but can infect Rab32 or BLOC-3 deficient mice (11).

*Salmonella* uses different host systems of post-translational modification to modify effectors towards its own benefit and survival (5). We were intrigued by the finding that GtgE contains a CaaX motif at its C-terminus (Fig 1A). This motif is recognized and targeted by host enzymes to introduce a hydrophobic isoprenoid modification to the Cysteine (7). This modification is required for those proteins that are targeted to membranes within the cell (7). We hypothesized that if GtgE is prenylated by the host, it would localize to membranes at which it can act on Rab32 more effectively, since activated Rab GTPases are in membranous compartments (10). To verify our hypothesis, we have generated a GtgE mutant that cannot be prenylated due to substitution of Cys^225^ for a Serine (GtgE^C225S^). We then used confocal microscopy to examine the intracellular localization of both YFP-GtgE and YFP-GtgE^C225S^ in HeLa cells. As shown in Fig 1B, YFP-GtgE localizes mainly in the perinuclear region showing a membrane-associated pattern. In contrast, YFP-GtgE^C225S^ is distributed throughout the cytoplasm and nucleus of the cell with no clear perinuclear or plasma membrane targeting. These results suggest that the CaaX motif is important to maintain GtgE on membrane structures. To confirm this finding, we performed a membrane fractionation of uninfected HeLa cell lysates after transfection with YFP-GtgE or YFP-GtgE^C225S^. As depicted in Fig 1C, YFP-GtgE^C225S^-expressing cell lysates showed a smaller proportion of the protein in the membrane enriched fraction compared to YFP-GtgE. This indicates that there is less amount of YFP-GtgE^C225S^ membrane-associated compared to YFP-GtgE.

**Fig 1.**
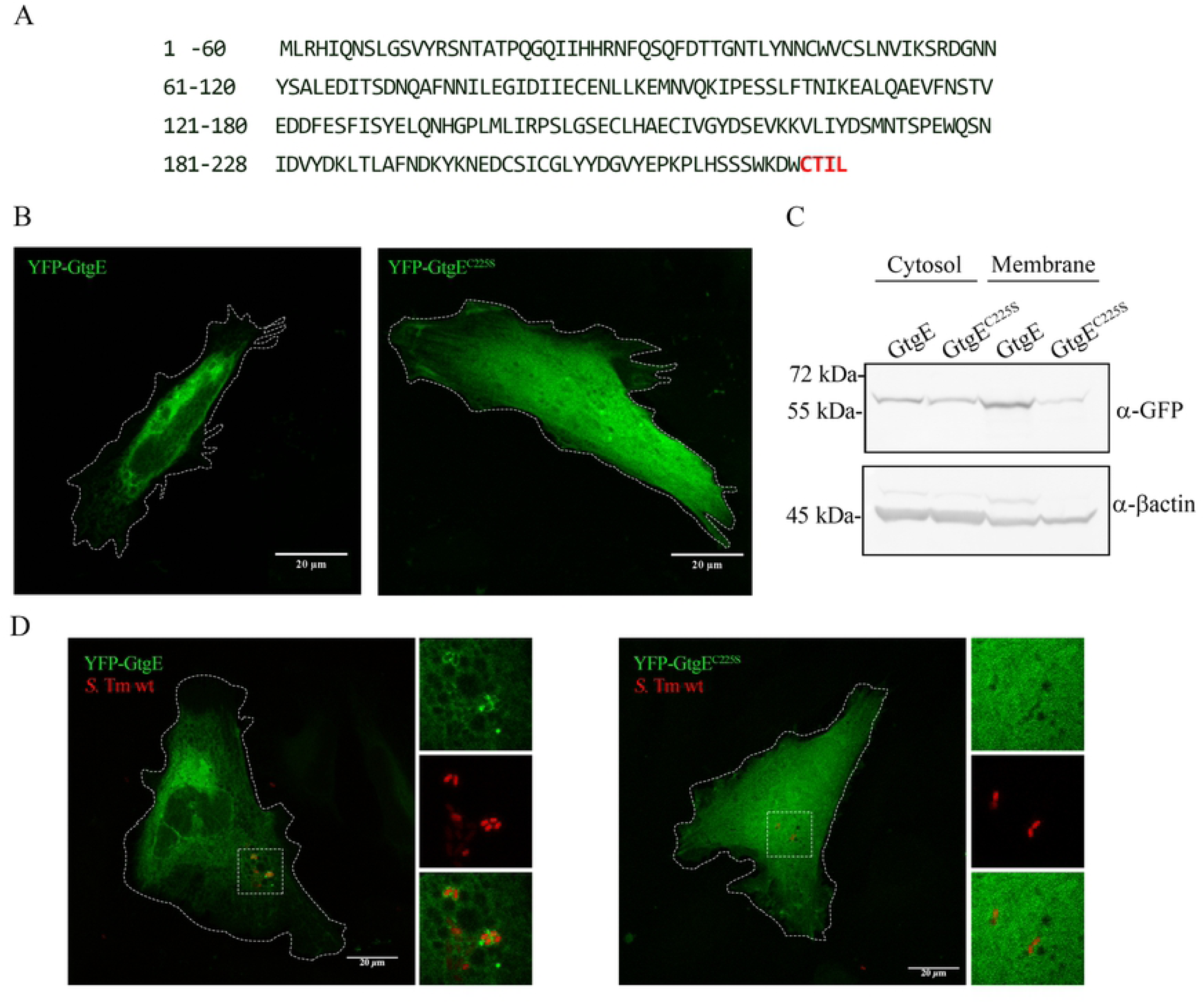
Localization of ectopically extressed YFP-GtgE depends on the predicted prenylated cysteine. (A) Sequence of GtgE shows a predicted CaaX motif at its C-terminus. (B) A substitution of GtgE Cys225 to Serine affects its intracellular distribution. HeLa cells were transfected to express YFP-GtgE or YFP-GtgEC225S (green) and fixed at 24 hours after transfection. Images show a perinuclear distribution of YFP-GtgE (left panel) and a nuclear/cytosolic distribution of YFP-GtgEC225S (right panel). Maximum-intensity projections of representative confocal Z-stacks are presented. (C) Prenylation of GtgE affects its subcellular distribution. Enriched cytosolic and membrane fractions were prepared from HeLa cells expressing YFP-GtgE or YFP-GtgEC225S. GtgE was detected with an anti-GFP antibody. β-actin was used as internal loading control. (D) GtgE localizes to the SCV. HeLa cells expressing YFP-GtgE or YFP-GtgEC225S were infected with *S*.Typhimurium mCherry for 2.5 hours. Images show YFP-GtgE decorating SCV (left panel) where no association to the SCV was observed in YFP-GtgEC225S expressing cells. Maximum-intensity projections of representative confocal Z-stacks are presented.

We have previously shown that Rab32 is efficiently recruited to the SCV of *S*. Typhimurium *ΔgtgEΔsopD2*, which leads to killing of the intracellular bacteria (11). It is therefore possible that cleavage of Rab32 by GtgE occurs on the SCV. If this is the case, GtgE should also localize to the SCV and this may be mediated by prenylation. To investigate the role of GtgE prenylation on its subcellular localization during infection, transiently transfected HeLa cells were infected with *S*. Typhimurium::mCherry wt. As shown in Fig 1D, YFP-GtgE^C225S^ and YFP-GtgE present the same differential distribution observed in uninfected cells. More importantly, YFP-GtgE but not YFP-GtgE^C225S^ can be found on the SCVs. This result suggests that prenylation is a necessary step for GtgE localization to the SCV. It is possible to hypothesize that Rab32 cleavage by GtgE may occur on the SCV. However, cleavage in other membrane compartments where Rab32 localizes (i.e. Golgi apparatus and Endoplasmic Reticulum) may still be possible.

### Translocated GtgE during infection localizes to membranes

In order to confirm that GtgE expressed during *Salmonella* infection is targeted to membranous compartments, we introduced a 3xFLAG sequence in the Asp^57^ position within a loop region of either GtgE or GtgE^C225S^. The resulting constructs (pSB065 and pSB069) were transformed into the *S*. Typhimurium *ΔgtgE* strain. Efficient translocation for both constructs was observed in translocation assays (Fig 2A) demonstrating that neither the triple FLAG insertion nor the C225S modification affect the translocation through the TTSS.

**Fig 2.**
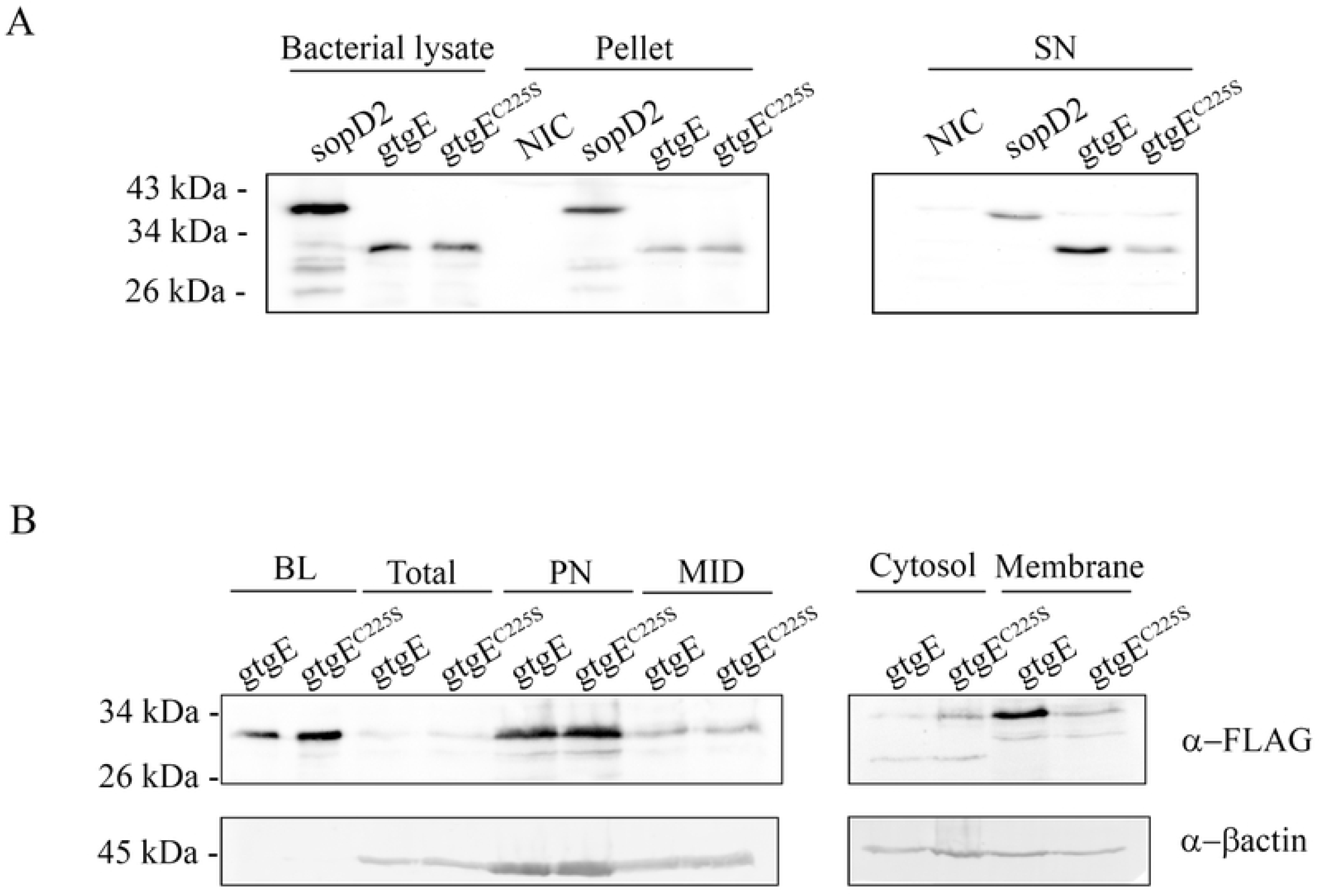
Translocated GtgE during infection is directed to membranes. (A) Translocation of 3xFLAG-tagged GtgE variants was determined. HeLa cells were infected with at MOI of 20 with *S*. Typhimurium *ΔsopD2* pSB4829 (SopD2-3xFLAG as positive control), *S*. Typhimurium *ΔgtgE* pSB065 (GtgE-3xFLAG) or S. Typhimurium *ΔgtgE* pSB069 (GtgE^C225S^-3xFLAG). After 5 hpi, cells were lysed and centrifuged at 10,000 x g for 10 minutes, the supernatant was filtered through a 0.2 μm filter to eliminate any remaining bacteria. Cell pellets and supernatant were used for Western blot, together with respective bacterial lysate from the inoculum used for infection, as positive control. NIC, non-infected control. Blots were developed with an anti-FLAG antibody. (B) Prenylation of translocated GtgE affects its subcellular distribution. HeLa cells were infected at MOI of 20 with *S*. Typhimurium *ΔgtgE* pSB065 (GtgE-3xFLAG) or *S*. Typhimurium *ΔgtgE* pSB069 (GtgE^C225S^-3xFLAG). After 4 hpi, cells were collected (Total sample), lysed and subjected to differential centrifugation, each pellet sample was used for Western blot analysis (PN, represents 500 x g pellet; MID, 10,000 x g pellet; membrane, 100,000 x g pellet), cytosol fraction was obtained by TCA precipitation of the resultant supernatant after 100,000 x g centrifugation. Detected with an anti-FLAG antibody. β-actin was used as internal loading control. BL indicates respective bacterial lysates from the inoculum used for infection.

HeLa cells were then infected with *S*. Typhimurium *ΔgtgE* harbouring pSB065 or pSB069. Four hours post-infection (hpi) the cells were harvested, and cytosolic and membrane fractions were collected. As shown in Fig 2B, the amount of GtgE^C225S^ detected in the membrane fraction is significantly lower compared to the membranous amount of GtgE. All together, these data suggest that translocated GtgE during cell infection is also prenylated to be targeted to membranous compartments.

### GtgE prenylation is required for efficient Rab32 cleavage

We and others have previously shown that GtgE specifically targets Rab29, Rab32 and Rab38, and that Rab32 cleavage is pivotal to counteract the Rab32-dependent antimicrobial pathway. We therefore investigated the effect of GtgE prenylation on Rab32 cleavage, generating *S*. Typhimurium strains (in wt and Δ*sopD2* backgrounds) expressing *gtgE*^*C225S*^. Crystallographic analysis showed that the C-terminus of GtgE is not a component of the catalytic site of the enzyme and that mutation of Cys^225^ does not affect GtgE protease activity (15). To assess the effect of C225S substitution on Rab32 cleavage, mouse *caspase 1*^*-/-*^ BMDM were infected with *S*. Typhimurium wt, *ΔgtgE, gtgE*^*C225S*^, *ΔsopD2, gtgE*^*C225S*^*ΔsopD2* or *ΔgtgEΔsopD2*. Cells were lysed and collected at 45, 70 and 150 minutes post-infection (mpi), and Rab32 cleavage was evaluated by Western-blot (Fig 3A). As expected, *ΔgtgE* strains do not show any Rab32 cleavage. In contrast, *S*. Typhimurium wt shows a clear proteolytic activity even at early time points (45 mpi). More interestingly, the prenylation defective mutant *gtgE*^*C225S*^ shows significantly less cleavage than *S*. Typhimurium wt at early time points, but cleavage was not significantly different at 150 mpi (Fig 3B). Therefore, even if GtgE prenylation is not strictly required for Rab32 cleavage, it does increase the effector proteolytic efficiency.

**Figure 3.**
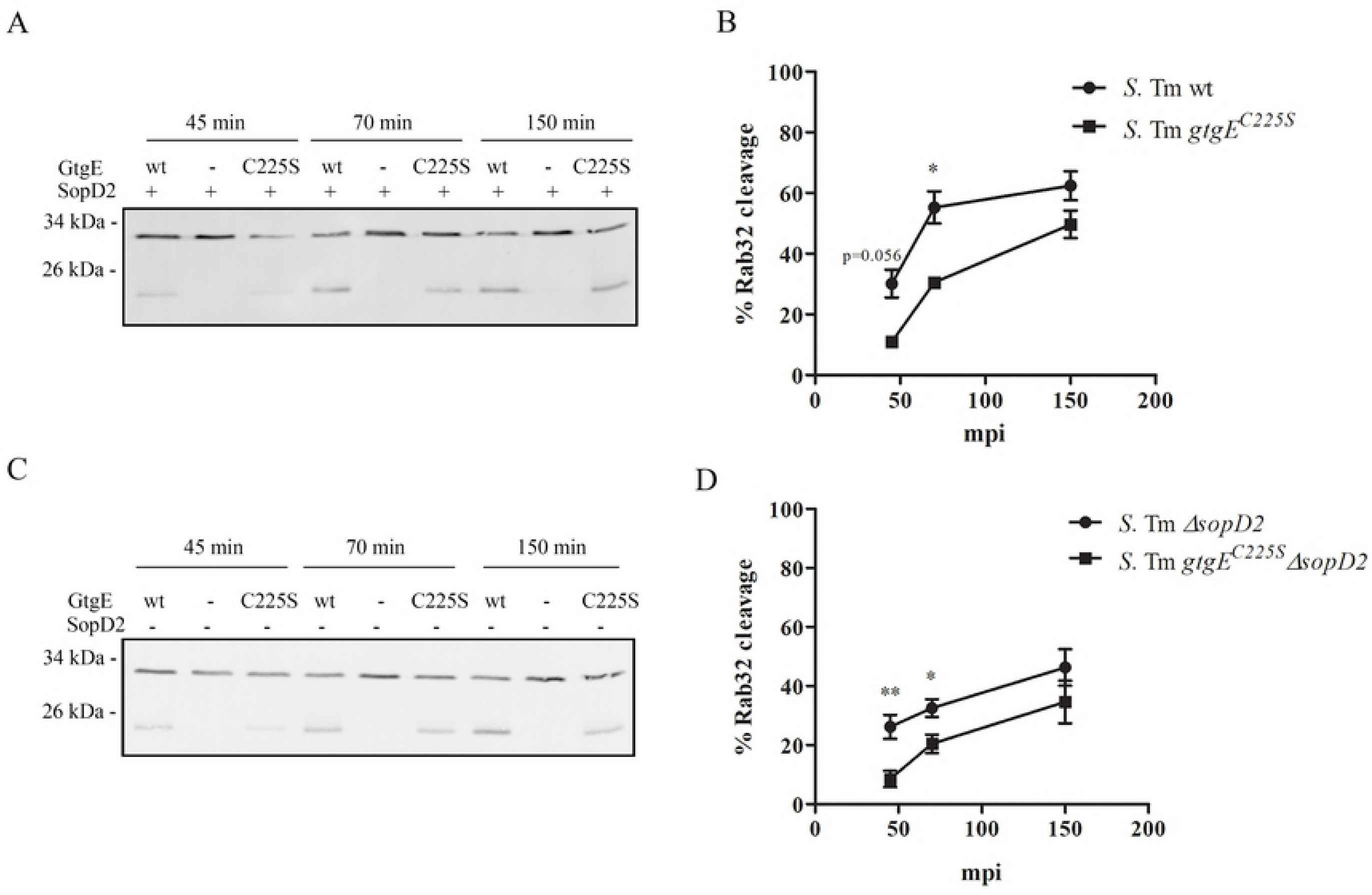
GtgE prenylation is important for Rab32 cleavage at the early stage of S. Typhimurium infection. (A) Mouse caspase 1 -/- BMDMs were infected with *S*. Typhimurium wt, *gtgE* and *gtgE*^*C225S*^. After 45, 70 and 150 mpi, infected cells were lysed and analysed by western blot with a mouse antibody against mouse Rab32.Images are representative western blot of Rab32 cleavage. (B) Rab32 cleavage quantification in BMDM infected with *S*. Typhimurium wt and *gtgE*^*C225S*^. The mean ± SEM of percentage of cleaved Rab32 in at least two independent experiments are shown. (C) Mouse caspase 1 -/- BMDMs were infected with *S*. Typhimurium *sopD2, ΔgtgEΔsopD2* and *gtgE*^*C225S*^*ΔsopD2*. After 45, 70 and 150 mpi, infected cells were lysed and analysed by western blot with a mouse antibody against mouse Rab32. (D) Rab32 cleavage quantification in BMDM infected with *S*. Typhimurium *ΔsopD2* and *gtgE*^*C225S*^*ΔsopD2* strains. The mean ± SEM of percentage of cleaved Rab32 in at least two independent experiments are shown.

It has been proposed that SopD2 could increase GtgE activity by elevating the amount of inactive, GDP bound, Rab32 (12). We therefore assessed Rab32 cleavage in a *ΔsopD2* background (Fig 3C). As shown in Fig 3D, a decrease in Rab32 cleavage was observed when GtgE^C225S^ was introduced in a Δ*sopD2* background, indicating that GtgE prenylation is important to ensure a quick effect on Rab32, independently of SopD2.

### GtgE prenylation is important for *Salmonella* virulence

Since GtgE prenylation seems to control the intracellular localization and is important for an efficient and fast cleavage of Rab32, we hypothesized that this post-translational modification would also have an impact in *Salmonella* intracellular survival. To evaluate this, we used a gentamicin protection assay to compare the intracellular survival of *S*. Typhimurium wt, *gtgE*^*C225S*^, *ΔgtgE ΔsopD2, gtgE*^*C225S*^*ΔsopD2* or *ΔgtgEΔsopD2* strains in mouse *caspase 1*^*-/-*^ BMDM (Fig 4A). As expected, and previously reported, strains lacking GtgE (*ΔgtgE* and *ΔgtgEΔsopD2*) show significantly less survival both at 5 and 24 hpi while no impairment was observed in a *ΔsopD2* strain (11, 12). Interestingly, the prenylation *gtgE*^*C225S*^ mutant shows a defective intracellular survival at 5 hpi both in wt and *ΔsopD2* background. This survival impairment, nevertheless, is no longer observed at 24 hpi. All together these results suggest that prenylation and therefore localization of GtgE plays a critical role at early stages of infection.

**Figure 4.**
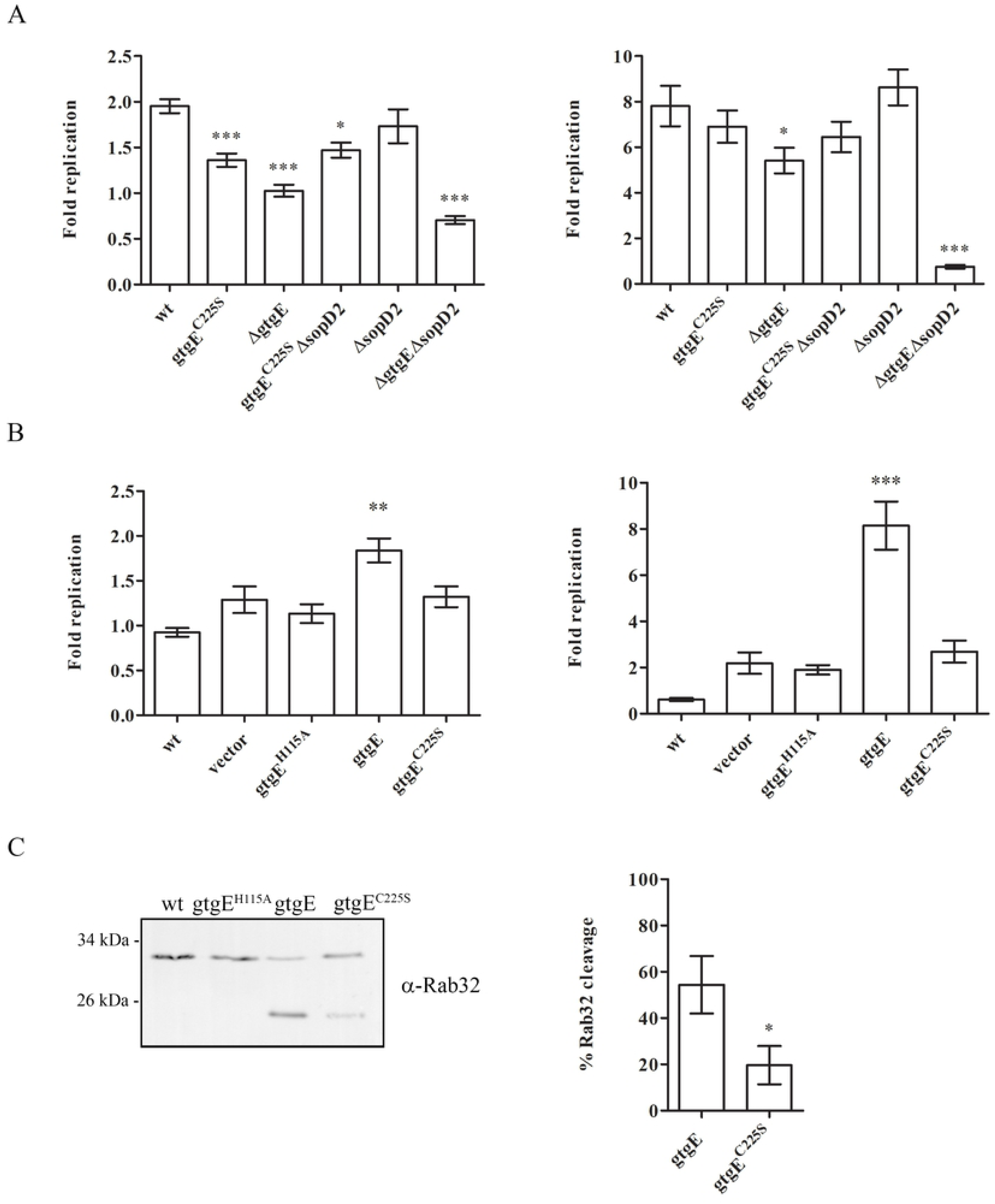
GtgE prenylation is important for macrophage intracellular survival. (A) Mouse BMDM caspase-/- cells were infected at MOI of 2 with *S*. Typhimurium wt, *S*. Typhimurium *gtgE*^*C225S*^, *S*. Typhimurium *ΔgtgE, S*. Typhimurium *ΔsopD2, S*. Typhimurium *gtgE*^*C225S*^*ΔsopD2* or S. Typhimurium ΔgtgE ΔsopD2. Cells were lysed at 1.5, 5 and 24 hpi and CFU were enumerated. Fold replication was calculated at5 hpi (left panel) and 24 hpi (right panel) versusinitial time point (1.5 hpi). Mean CFU ± SEM of at least three independent experiments are presented. P values were assessed by one-way ANOVA with Dunnett’s posttest. (B) Mouse BMDM caspase-/- cells were infected at MOI of 10 with *S*. Typhi wt, *S*. Typhi with pSB4004 (vector), *S*. Typhi pSB4135 (*gtgE*^*H115A*^), *S*. Typhi pSB065 (*gtgE*) or *S*. Typhi pSB069 (*gtgE*^*C225S*^). Cells were lysed at 1.5, 5 and 24 hpi and CFU were enumerated. Fold replication was calculated at5 hpi (left panel) and 24 hpi (right panel) versusinitial time point (1.5 hpi). Mean CFU ± SEM of at least three independent experiments are presented. P values were assessed by one-way ANOVA with Dunnett’s posttest. (C) Rab32 cleavage was examined by Western blot in mouse BMDM caspase-/- cells after 2.5 hpi infection with a MOI of 10 of *S*. Typhi wt, *S*. Typhi pSB4135 (*gtgE*^*H115A*^), *S*. Typhi pSB065 (*gtgE*) or *S*. Typhi pSB069 (*gtgE*^*C225S*^). Quantification of Rab32 cleavage was performed in ImageJ, average values ± SEM of at least three experiments are presented. Statistical differences were assessed by means of unpaired Student’s t-test, with *p<0.05.

To further investigate the importance of GtgE prenylation during infection, we decided to study the effect of GtgE^C225S^ in *S*. Typhi (Fig 4B). *S*. Typhi does not express *gtgE* or *sopD2*, and therefore is sensitive to the Rab32 antimicrobial pathway in mouse macrophages. However, we have demonstrated that the sole expression of GtgE is enough to increase *S*. Typhi survival in mouse macrophage (11). Therefore, *S*. Typhi represents a perfect model to individually study the contribution of GtgE to *Salmonella* virulence. *S*. Typhi was transformed with plasmids encoding *gtgE, gtgE*^*C225S*^ or *gtgE*^*H115A*^, a catalytically inactive form of GtgE (12) and used to infect mouse *caspase 1*^*-/-*^ BMDM. As previously reported, expression of *gtgE* conferred a significant increase of *S*. Typhi survival both at early and late time points (11). However, expression of *gtgE*^*C225S*^ does not provide any advantage for intracellular replication at any time point. As for *S*. Typhimurium *gtgE*^*C225S*^, a delay in Rab32 cleavage was also observed in *S*. Typhi pGtgE^C225S^ (Fig 3C), showing again that prenylation and localization are important for an efficient enzymatic activity. Interestingly, in the *S*. Typhi model, these differences were maintained at 24 hpi, and the behaviour of *S*. Typhi with GtgE^C225S^ was virtually the same as expressing a catalytically inactive form of GtgE, suggesting that effective localization of GtgE is as important as a catalytically active enzyme to efficiently counteract Rab32 in *S*. Typhi. Differences in survival observed at 24 hpi between *S*. Typhimurium *gtgE*^*C225S*^ and *S*. Typhi expressing *gtgE*^*C225S*^ may be the result of other effectors or ways to respond to the antimicrobial environment of the SCV imposed by Rab32.

Finally, to investigate the contribution of GtgE prenylation to *Salmonella* virulence *in vivo*, C57Bl/6 *caspase 1*^*-/-*^ mice were intraperitoneally infected with *S*. Typhimurium *ΔsopD2, gtgE*^*C225S*^*ΔsopD2* or *ΔgtgEΔsopD2*. In this case, we decided to work only in a *ΔsopD2* background in order to exclude the contribution of this effector to the Rab32 pathway and concentrate only upon the role of GtgE. The ability of the strains to establish a systemic infection was assessed by CFU recovery from spleens at 4 days post-infection (dpi) (Fig 5). As expected, *ΔsopD2* was recovered at higher load number (median ≈ 6 x 10^6^) compared to *ΔgtgEΔsopD2* (median ≈ 1 x 10^4^). Interestingly, *gtgE*^*C225S*^*ΔsopD2* presented an intermediate phenotype, showing a significantly less bacterial load than *ΔsopD2* (median ≈ 2 x 10^5^). This indicates that prenylation of GtgE confers a virulence advantage for *S*. Typhimurium to establishing a systemic infection. Taken together, these results suggest that prenylation of GtgE has an important role at cellular and systemic levels in *S*. Typhimurium infection.

**Figure 5.**
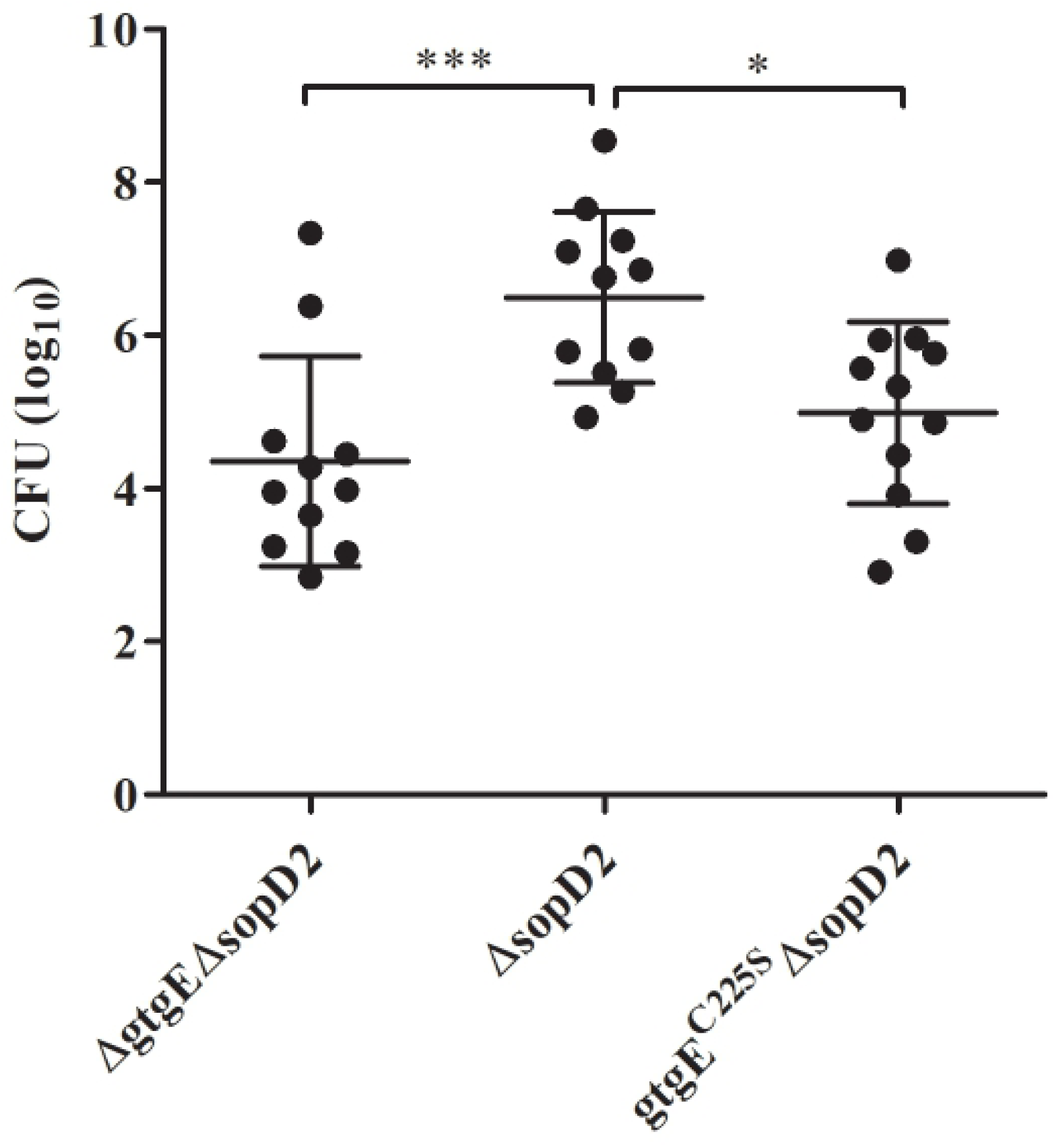
GtgE prenylation contributes to S. Typhimurium virulence. C57BL/6 caspase 1−/−mice were intraperitoneally inoculated with 1 x 103 CFUs of *S*. Typhimurium *ΔsopD2, S*.Typhimurium *gtgE*^*C225S*^*ΔsopD2* or *S*. Typhimurium *ΔgtgEΔsopD2*, and 4 days post-infection, number of bacteria recovered from the spleen of infected mice were enumerated. Data from two independent experiments are presented, with median and SD values. Indicated p values were determined by one-way ANOVA test with Dunnett’s posttest.

## Discussion

Manipulating host activities through translocated effectors is crucial for intracellular survival in bacterial pathogens. Host post-translational modifications of bacterial effectors have been extensively reported, such as ubiquitination, phosphorylation or lipidation (16); yet, prenylation remains understudied. Al-Quadan et al. proposed a list of potential prenylation targets from different bacterial pathogens with a C-terminal CXXX motif. One of these proteins is GtgE, which contains a CTIL sequence at its C-terminus (17). Moreover, Krzysiak et al. showed that a CTIL peptide can be farnesylated (18). These results suggest that GtgE could be subjected to lipidation in the host cell during infection, and that this modification could be important for its function. In this work we studied GtgE localization upon prenylation of the cysteine in its CTIL C-terminal sequence. We found that replacing Cys^225^ by Ser changes GtgE distribution in transfected HeLa from the perinuclear region, with a membrane-associated pattern to a cytosolic and nuclear localization, as evidenced by confocal microscopy. This same observation was found in prenylated effectors AnkB from *Legionella pneumophila* and SifA from *S*. Typhimurium when an analogous mutation was introduced in their protein sequences (19-21). We then confirmed the differences in localization by Western-blot and subcellular fractionation. Moreover, we were able to localize YFP-GtgE to the SCV during infection of transfected HeLa cells, whereas no association to the SCV was observed in YFP-GtgE^C225S^ expressing cells. Prenylated AnkB was also observed to be directed to the *Legionella*-containing vacuole, which promotes the evasion of the endosomal-lysosomal pathway (19, 20). Localization of GtgE to the SCV could suggest that this is the site where Rab32 cleavage takes place.

Although results from our studies with HeLa transfected cells showed striking differences between GtgE^C225S^ and GtgE localization, we sought to understand if this post-translational modification was indeed occurring during *Salmonella* infection. Because of technical limitations found for microscopic visualization of GtgE during infection, we decided to do a fractionation assay of HeLa infected cells where *S*. Typhimurium would express GtgE^C225S^-3xFLAG or GtgE-3xFLAG. These experiments showed less abundance of GtgE^C225S^-3xFLAG in membrane fractions in comparison with GtgE-3xFLAG, indicating that prenylation does affects localization of GtgE during infection.

GtgE functions as a protease that targets Rab29, Rab38 and Rab32 (13). It was previously shown by us that GtgE together with SopD2 act together to inactivate Rab32, playing a role in the survival of intracellular *Salmonella* (11). Rab32 in its inactive GDP-bound form would be solubilized to the cytosol by interaction with GDI. In this context, GtgE cannot cleave Rab32, since GDI and GtgE interact with similar binding sites on Rab32, and GDI-Rab interactions have high affinity (22). Therefore, GtgE would need to access Rab32 membrane-bound forms. We observed that Rab32 cleavage was less effective in GtgE^C225S^ expressing *S*. Typhimurium and *S*. Typhi strains. Our results indicate that, even though GtgE^C225S^ can still cleave Rab32, localizing GtgE to membranes will provide a faster and more effective response to the pathway. This could explain the defect in survival at early time points of infection in *S*. Typhimurium and in *S*. Typhi expressing GtgE^C225S^ variants. In addition, Rab32 cleavage was not affected in Δ*sopD2* genetic background strains. This seems to contradict Watchel et al, who proposed that GtgE only targets inactive Rab GTPases (22). However, it is possible to reconciliate this discrepancy if we hypothesize that there is another Rab32 GAP in the process, either from host or bacterial origin.

Surprisingly, the use of *S*. Typhi expressing GtgE as model, has highlighted important differences with *S*. Typhimurium. Rab32 cleavage was found to be slower in *S*. Typhi than in *S*. Typhimurium and, more importantly, *S*. Typhimurium *gtgE*^*C225S*^ can recover from Rab32 cleavage defect in mutants at later time points, but *S*. Typhi cannot. These differences in survival between the serovars may be explained by other mechanisms present in *S*. Typhimurium, but absent in *S*. Typhi, to cope with the SCV environment induced by Rab32. For example, it was recently demonstrated that Rab32 assists in the delivery of itaconate acid from mitochondria to the SCV. Interestingly, *S*. Typhimurium expresses genes involved in itaconate acid degradation, while *S*. Typhi does not (14). It is then possible that *S*. Typhimurium can survive longer in presence of an active Rab32 pathway and a slower action of the non-prenylated GtgE has less impact on *S*. Typhimurium survival than *S*. Typhi. In any case, the importance of a fast neutralization of the Rab32 pathway for virulence is demonstrated by our experiments in a mouse model. *S*. Typhimurium *gtgE*^*C225S*^ shows a decreased ability to colonize mice, suggesting that even a delay in Rab32 cleavage results in reduced bacterial virulence. *S*. Typhimurium has evolved mechanisms to subvert host activities through action of translocated effectors that allow its successful intracellular survival. Even further, like many other bacterial pathogens, *S*. Typhimurium uses host machineries to introduce post-translational modifications in its effectors, ensuring an effective function within the cell (21, 23, 24). In this study we showed that prenylation of GtgE is essential to its cellular localization, to efficiently target Rab32 and, consequently, to successfully establish infection.

**This paper is part of Stefania Spanò’s scientific legacy and this would have not been possible without her intelligence, vision and persistence. A dreadful destiny has snatched her from us too early, but her discoveries and ideas are living and flourishing**.

## Materials and methods

### Strains and growth conditions, plasmids and primers

All strains, plasmids and primers used in this work are listed in Table 1. *S*. Typhimurium and *S*. Typhi strains were maintained on LB agar. When appropriate, antibiotic was added at the following concentrations, streptomycin, 50 µg mL^-1^; kanamycin, 30 µg mL^-1^, tetracycline 12.5 µg mL^-1^. Liquid cultures were grown on LB broth or LB broth with 0.3 M NaCl for induction of SPI-1 TTSS genes.

**Table 1.**
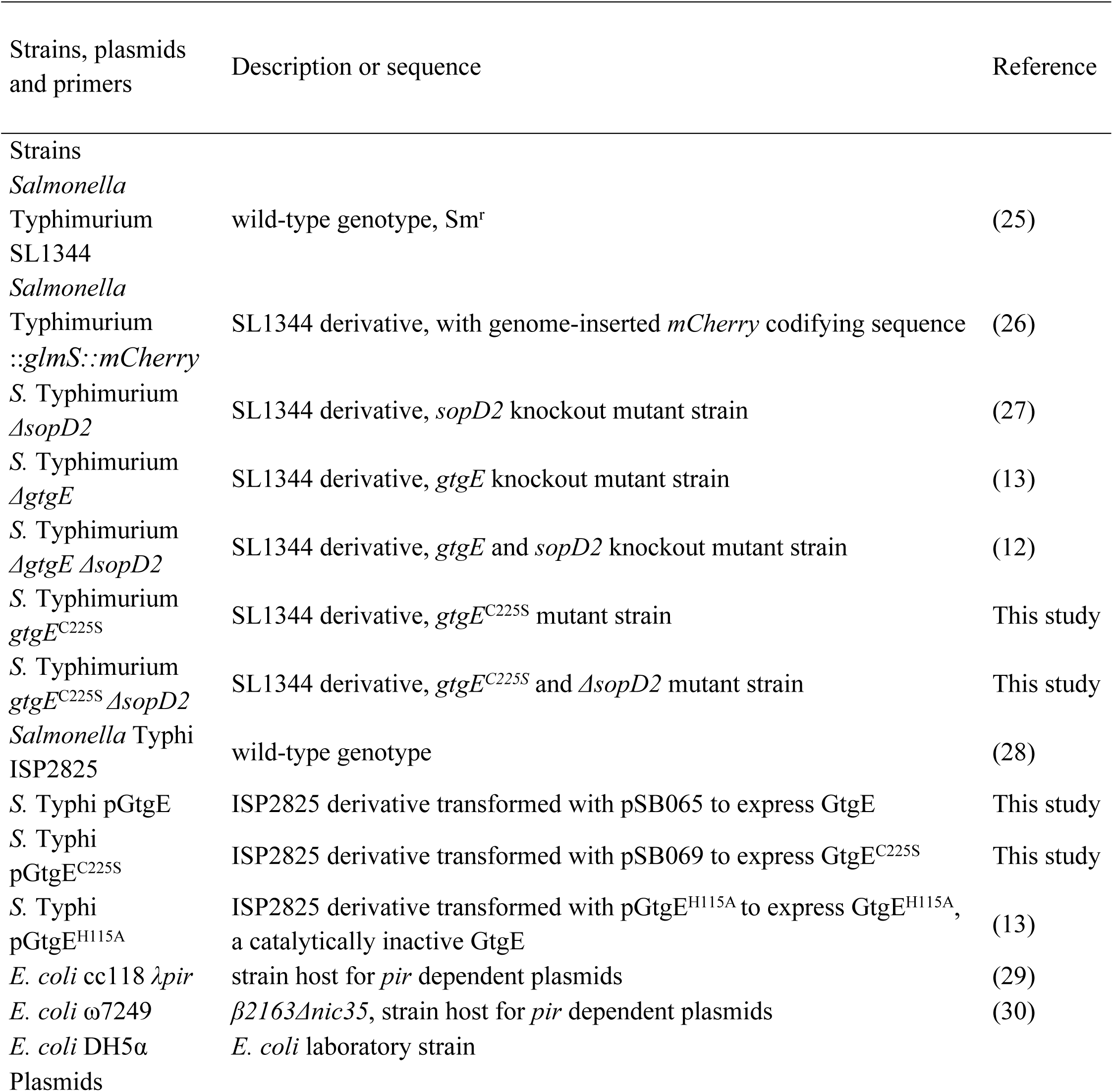
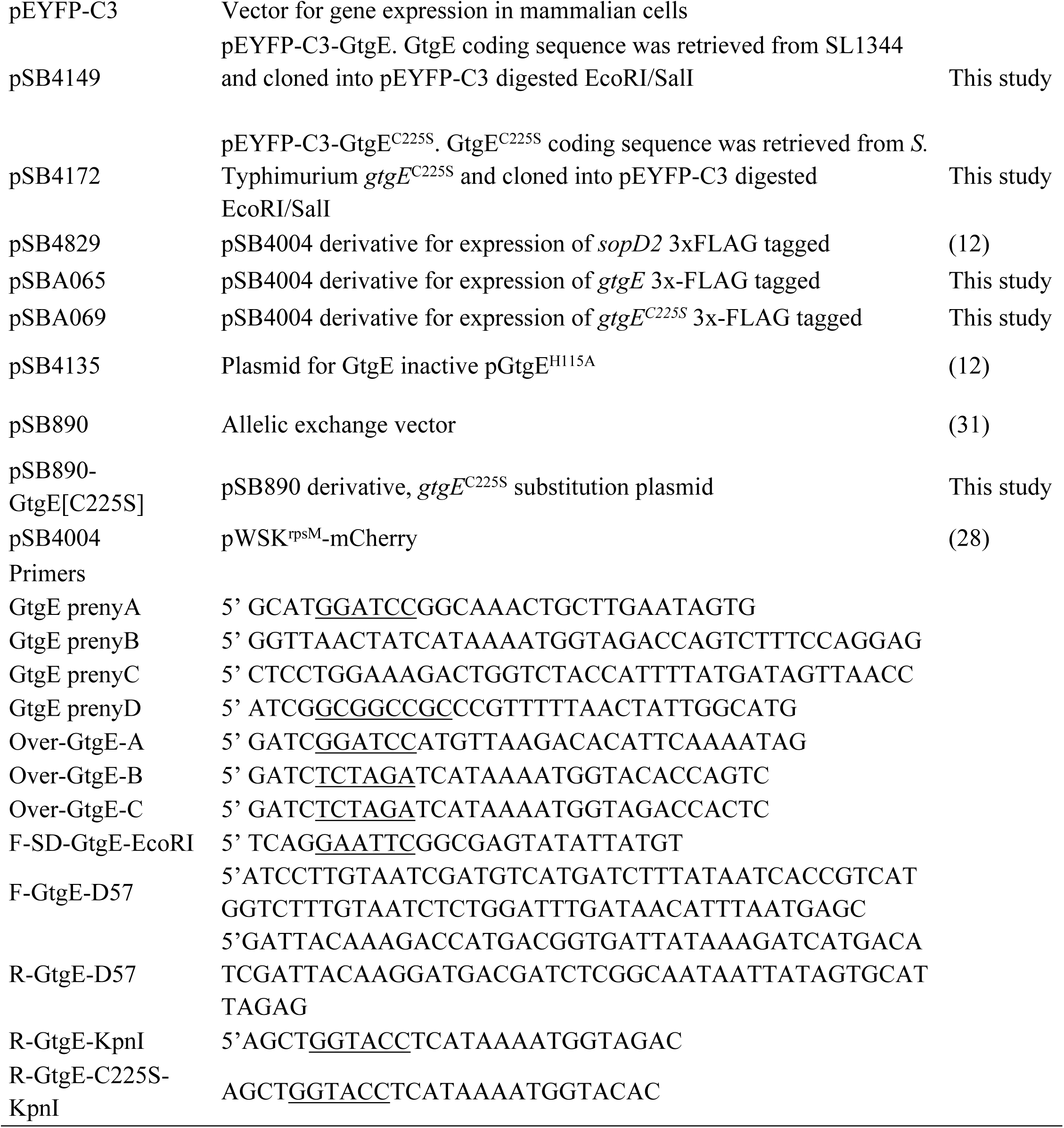
Strain, plasmids and primers used in this study.

### Cell Lines

HeLa and L929 were grown in Dulbecco’s Modified Eagle’s Medium (DMEM high glucose with glutamax, Gibco) supplemented with 10% Fetal Bovine Serum (FBS, Gibco) and cultured in 10 cm tissue culture (Fisher) plastic plates at 37°C, 5% CO_2_.

### Isolation of BMDM

Mouse *Caspase-1*^*-/-*^ BMDMs were isolated as previously described (32). Cells were cultured on RPMI 1640 medium (Gibco) supplemented with 10% FBS, 2 mM glutamine (Gibco) and 20% of L929 cells supernatant (BMDM medium), for 6 to 9 days before use for infection assays. Fresh BMDM medium was first replaced 3 days after plating and then every 2 days.

### Plasmid constructs generation and DNA manipulation Introduction of C225S substitution in GtgE

A point mutation was introduced at 674 nt position on *gtgE* in *S*. Typhimurium wt and *S*. Typhimurium *ΔsopD2* by allelic exchange as previously described (31). Briefly, the mutation was introduced by amplifying *gtgE* with the primer pairs GtgE-prenyA and GtgE-prenyB, and GtgE-prenyC and GtgE-prenyD by overlapping PCR. Amplicon was purified, double digested with BamHI/NotI (NEB) and ligated to BamHI/NotI digested pSB890 (31) to generate pSB890-GtgE[C225S] construct. pSB890-GtgE[C225S] was electroporated into *Escherichia coli* cc118 *λpir* strain and transformants were selected by tetracycline resistance, and correct sequence of the construct was corroborated. Purified pSB890-GtgE[C225S] was then transferred to *E. coli* ω7249 strain by electroporation and transformants were selected by tetracycline resistance on a LB plate supplemented with 50 µg mL^-1^ of diaminopimelate (DAP, Sigma-Aldrich) (30). An *E. coli* clone harbouring the pSB890-GtgE[C225S] with the correct inserted sequence was selected and used as donor strain in matting experiments, including *S*. Typhimurium wt or *S*. Typhimurium *ΔsopD2* as recipient strain. Exoconjugants were selected by tetracycline resistance and grown in 20 mL of LB broth at 30°C for 8 hours and counter selected for second recombination events on LB agar plates supplemented with 10% w/v sucrose. Clones that contained the point mutation were confirmed by PCR and AccI restriction enzyme digestion. Mutants were finally confirmed by sequencing.

### Overexpression constructs

For overexpression assays in eukaryotic cells, *gtgE* and *gtgE*^*C225S*^ were amplified with primers pair Over-GtgE-A and Over-GtgE-B, and Over-GtgE-A and Over-GtgE-C, respectively, from *S*. Typhimurium wt and *S*. Typhimurium *gtgE*^*C225S*^ respective genomes, digested with BamHI and XbaI and cloned into BamHI/XbaI digested pEYFP-C3 plasmid, to generate pSB4149 and pSB4172. Constructs were transformed into *E. coli* DH5α electrocompetent cells and selected by kanamycin resistance. Correct inserts were confirmed by sequencing.

### Insertion of 3xFLAG tag into GtgE and GtgE^C225S^

A 3xFLAG tag was introduced at Asp^57^ position of GtgE by overlapping PCR with primers F-SD-GtgE-EcoRI, F-GtgE-D57, R-GtgE-D57, R-GtgE-KpnI (for *gtgE* wt sequence) or R-GtgE-C225S-KpnI (for *gtgEC225S* sequence). Amplicons were digested with EcoRI/KpnI and cloned into EcoRI/KpnI digested pSB4004, to generate pSB065 (for *gtgE*-3xFLAG) and pSB069 (for *gtgE*^*C225S*^-3xFLAG). Constructs were transformed into *E. coli* DH5α electrocompetent cells and selected by kanamycin resistance. Correct inserts were confirmed by sequencing and then electroporated into *S*. Typhimurium or *S*. Typhi strains.

### Intracellular localization of GtgE

HeLa cells were seeded on coverglasses (#1 Thermo Scientific) in 24 well plates at a density of 2 x 10^4^ cells and grown overnight. PEI (60 μg mL^-1^, Sigma-Aldrich) or 1 μg of DNA (pSB4149 or pSB4172) was added to DMEM medium without FBS and incubated for 10 minutes at room temperature, then mixed in 1:1 ratio and incubated for additional 20 minutes at room temperature. HeLa cells were washed once with PBS and transfected with 100 μL of the DNA/PEI mix, 500 μL of growth media was added and cells were incubated for 24 hours, before fixation with 4% PFA and further mounting. Transfected cells were also infected with MOI of 20 with *S*. Typhimurium wt chromosomally expressing mCherry, following the gentamycin protection assay detailed below. After 2.5 hpi, cells were washed, fixed and samples mounted with ProLong Diamond antifading agent (ThermoFisher). Samples were visualized with a Zeiss LSM880 microscope, z-stack sections were taken, and maximum projections were obtained. Experiments were repeated twice with duplicates.

### Fractionation of cellular compartments

Fractionation of cellular compartments was performed by multi-step centrifugation according to Gomes et al. with minor modifications (33). HeLa cells were transfected to express either YFP-GtgE wt or YFP-GtgE^C225S^ with plasmids pSB4149 and pSB4172 respectively. Cells were seeded at a density of 5 x 10^5^ cells in 10 cm culture dishes and grown overnight. The next morning, 12 μg of plasmid DNA (pSB4149 and pSB4172) or PEI (60 μg/mL) were added to DMEM medium for 10 minutes at RT, then mixed in 1:1 ratio and incubated for additional 20 minutes. Meanwhile, the cells were washed once with PBS. HeLa cells were transfected with 1 mL of the DNA/PEI mix and incubated for 24 hours. In addition, fractionation was performed on infected HeLa cells with *S*. Typhimurium harboring either pSB065 or pSB069. Cells were seeded at 3 x 10^6^ cell density in 15 cm culture dishes and grown overnight. Next, HeLa cells were infected with a MOI of 20 with either *S*. Typhimurium strains, following the gentamycin protection assay for 4 hpi (see below). For both cases, transfected or infected cells were washed twice with warm PBS, harvested by trypsinization, spun down at 750 x g for 5 minutes, washed once with PBS and suspended in Homogenization Buffer (HB: 250 mM sucrose, Sigma-Aldrich; 10 mM HEPES pH 7.4, Sigma-Aldrich; 1 mM EDTA, Sigma-Aldrich, 2 mM PMSF, Roche). Homogenization was performed by a single freeze/thaw cycle, followed by repeated passage of the cellular material through a 25-gauge needle. Twenty μL of this suspension were set aside for Western blot analysis, as a control of GtgE or GtgE^C225S^ expression (referred as Total sample). Removal of cellular debris was achieved by low speed centrifugation at 500 x g for 10 minutes at 4°C. Subsequently, the supernatant was spun down at 100,000 x g for 1 hour at 4°C; however, infected HeLa were subjected to centrifugation at 10,000 x g for 10 minutes at 4°C prior to the ultracentrifugation step. Each of the fractionation steps including pellet after 500 x g (PN sample), 10,000 x g (MID sample), 100,000 x g (membrane fraction) centrifugations were suspended in 100 μL of 2X loading buffer (4% SDS, 10% β-mercaptoethanol, Sigma-Aldrich, 20% (v/v) glycerol, Sigma-Aldrich, 125 mM TRIS, Sigma-Aldrich and 0.004% bromophenol blue, Sigma-Aldrich), whilst the ultracentrifugation supernatant (cytosol fraction) was further processed in pursuance of TCA precipitation, and protein pellets were suspended in 200 μL of 2x Loading Buffer. Fractions were used directly for SDS-PAGE and immunoblotting. Transfection and infection experiments were repeated at least three times.

### Translocation

Translocation assay of TTSS effectors into the host cells was performed as previously described (11). Briefly, HeLa cells were seeded at 5 x 10^5^ in 6-well plate and grown overnight. The following day, cells were washed 3 times with HBSS, infected with *S*. Typhimurium wild-type, *S*. Typhimurium *ΔgtgE* pSB065 (encoding GtgE-3xFLAG), *S*. Typhimurium *ΔgtgE* pSB069 (encoding GtgE^C225S^-3xFLAG) and *S*. Typhimurium *ΔsopD2* pSB4829 (encoding SopD2-3xFLAG, as a positive control) at a MOI 20 for 5 hours, following the gentamycin protection assay. Proteinase K (30 µg/ml) suspended in HBSS was added 15 minutes prior the infection end point; incubated at 37°C to detach the cells and inhibited by addition of 2 mM PMSF solution in HBSS. Cells were harvested and spun down at 300 *x* g, 5 minutes. Subsequently, the cellular pellet was dissolved in 100 μL of the ice-cold lysing buffer (0.2% Triton X-100 (Sigma-Aldrich) in PBS; 2 mM PMSF) and centrifuged at 20,000 *x g*, 5 minutes at 4°C. Finally, the supernatant was collected and filtered through a Millipore® Millex® LG Syringe Filter with Hydrophilic LCR PTFE Membrane (0.2 µm) by centrifugation at 20,000 *x g*, 30 s at 4°C. Pellet and filtered fraction were mixed with 100 μL of 2X loading buffer or 20 μL of 5X loading buffer accordingly and analysed by SDS-PAGE and immunoblotting.

### Gentamycin protection assay

Bacterial intracellular replication assays were conducted as previously described (12). Briefly, mouse *caspase-1*^*-/-*^ BMDMs were seeded at a density of 6 x 10^4^/well in 24-well plates. *S*. Typhimurium or *S*. Typhi strains were diluted 1:20 on LB with 0.3 M NaCl from an overnight culture and grown until OD_600_ of 0.9 was reached. Prior to infection, BMDMs were washed twice with Hank’s balance salt solution (HBSS, Gibco) and infected with a MOI of 2 for *S*. Typhimurium or MOI of 10 for *S*. Typhi strains, prepared on HBSS. One-hour post-infection, cells were washed twice with HBSS and incubated with BMDM medium supplemented with 100 µg mL^-1^ gentamicin for 30 min. Cells due to be lysed at 1.5 h post-infection were washed twice with 0.5 ml and 1 ml of PBS, respectively, and lysed in 1 ml of 0.1% sodium deoxycholate (DOC, Sigma-Aldrich) in PBS by incubating at RT for 10 minutes and pipetting several times. For cells to be lysed at later time points, DMEM with 100 μg mL^-1^ gentamicin was replaced with fresh DMEM containing 5 µg mL^-1^ gentamicin to avoid cycles of reinfection. At the time of lysis (5 and 24 hpi), cells were incubated with growth medium containing 100 µg mL^-1^ gentamicin for 20 minutes and lysed with 0.1% DOC. After lysis, the number of intracellular bacteria was calculated by CFU determination. Experiments were repeated at least three times, significance was determined by means of one-way ANOVA and Dunnett’s post-test.

### Rab32 cleavage

Mouse *caspase-1*^*-/-*^ BMDMs were seeded at a density of 8 x 10^4^ cells in 24 well plates, and the next day were infected with *S*. Typhimurium strains at a MOI of 2. At 45, 70 and 150 minutes post-infection, BMDMs were washed with PBS and lysed in 20 μl of Laemmli loading buffer. Cell lysates were transferred to microcentrifuge tubes and boiled for 5 minutes to be analysed by Western blotting. Experiments were repeated at least two times, quantification of Rab32 cleavage was performed in ImageJ, significance was assessed by means of unpaired Student’s t test.

### Western blotting

Protein samples were separated by 12.5% SDS-PAGE, transferred to PVDF membranes (Immobilon; Millipore) with a semi-dry system (Bio-Rad) and then subjected to immunochemical detection. Primary antibodies were prepared in 2.5% skimmed milk and TBS-0.05% Tween (TBST), at the following dilutions: rabbit anti-GFP dilution (1:3,000, Abcam ab6556); rabbit anti-β-actin (1:1,000, Cell Signalling 4967), mouse anti-mouse Rab32 (1:1,000, Santa Cruz sc-390178); mouse monoclonal anti-FLAG M2 (1:5,000, Sigma F3165). Blocked membranes were incubated overnight at 4°C with the respective primary antibody. Next, the corresponding secondary antibody prepared at the following dilutions were added and incubated for 1h at room temperature, donkey anti-rabbit-800 (1:15,000, LI-COR 926-32213), donkey anti-mouse-800 (1:15,000, LI-COR 926-32212), goat anti-mouse HRP (1:10,000, Sigma A4416). Blots were developed by either ECL-detection method (Thermo Fisher) or by the Odyssey infrared imaging system (LI-COR Biosciences).

### Animal experiments

Groups of 6 C57Bl/6 *caspase-1*^*-/-*^ mice of 8-12 weeks were intraperitoneally inoculated with a dose of approximately 10^3^ CFU of *S*. Typhimurium Δ*sopD2, S*. Typhimurium Δ*gtgE*Δ*sopD2* and *S*. Typhimurium *gtgE*^*C225S*^Δ*sopD2* strains grown on LB 0.3 M NaCl until OD600 of 0.9. After 4 days post-infection, mice were sacrificed, and spleens were harvested and homogenized in 3 mL of PBS containing 0.05% DOC for CFU determination. The experiment was independently repeated twice, and statistical analysis was performed by means of one-way ANOVA and Dunnett’s post-test.

### Ethical Statement

All animal research was carried out within the Medical Research Facility, University of Aberdeen, in compliance with the conditions required by the UK Government Home Office as described under the Animals (Scientific Procedures) Act 1986 including European Directive 2010/63/EU.

These conditions include but are not limited to the sourcing of animals, animal care and welfare, health monitoring, the provision of veterinary treatment, housing and environmental conditions, the performance of all procedures, including humane killing. These conditions also include training, supervision and competence assessment requirements and the establishment and maintenance of a local Animal Welfare and Ethical Review Body. All animals used are bred and supplied by Home Office approved suppliers, regularly health screened and housed under Specific Pathogen Free (SPF) conditions (Project Licence number: 70/8073). All members of the animal care staff undergo a University approved training scheme and most hold their own personal licences.

The Medical Research Facility is a modern purpose-built facility and is subject to regular and on-going inspections by the Home Office.

## Author contribution

MB and SS designed and supervised the study. NC, HY and KP designed, performed and analysed the experiments. NC wrote the manuscript with MB supervision. All authors revised the manuscript and contributed with discussion and interpretation of the results.

## Acknowledgements

We are very grateful to Leigh Knodler for her generous gift of P22 phages from a *S*. Typhimurium *glmS::Cm::mCherry* strain. We would also like to thank to Prof Heather Wilson and Prof Gordon Dougan for their valuable and constructive suggestions during the manuscript drafting.

